# Integrated Representations of Threat and Controllability in the Lateral Frontal Pole

**DOI:** 10.1101/2025.08.24.671754

**Authors:** Joanne E. Stasiak, Jingyi Wang, Neil M. Dundon, Elizabeth J. Rizor, Christina M. Villanueva, Parker L. Barandon, Scott T. Grafton, Regina C. Lapate

## Abstract

Emotional processing is ubiquitous in everyday life, informing goal pursuit not only in response to current demands, but also in anticipation of future outcomes. Lateral prefrontal (LPFC) function supports cognitive control, and emerging evidence suggests a unique role for its anterior-most region—the lateral frontal pole (FPl)—in integrating putatively amygdala-originated emotion signals with goal information. However, whether these organizational properties of LPFC are expressed during the anticipation of future threat remains unknown. Here, we used FIR modeling and pattern similarity analysis to examine dynamic engagement and representational properties of distinct LPFC regions during threat anticipation requiring goal-directed action. Healthy participants (n=67) were scanned during a threat-of-shock paradigm consisting of a prolonged (18s) countdown to possible shock administration. Threat unpleasantness and controllability were manipulated orthogonally: in controllable trials, participants could avoid an unpleasant or mild shock by making a successful time-sensitive response; in uncontrollable trials, shocks were administered regardless of performance. LPFC robustly coded for anticipated threat unpleasantness, with FPl showing the strongest modulation by threat unpleasantness and controllability relative to caudal and mid-LPFC regions. Caudal and mid-LPFC maintained independent representations of threat unpleasantness and controllability. In contrast, FPl held conjunctive threat-and-controllability representations, which were associated with successful motor performance following anticipation of unpleasant shocks. Stronger conjunctive FPl representations were also associated with greater inverse amygdala–FPl coupling. Together, these findings provide insight into LPFC organization under naturalistic emotional challenges and highlight a key role for FPl in integrating affective and control-related information during threat anticipation to support goal-directed action.

**Significance Statement:** Anticipating emotionally-charged events—such as a painful outcome we may or may not be able to avoid—requires integrating emotion and control to guide behavior. However, the neural mechanisms through which emotional states influence goal-directed behavior in naturalistic, anticipatory emotional contexts remain unclear. Using a threat-of-shock paradigm and multivariate analyses we show that the anterior-most region of the lateral prefrontal cortex (LPFC)—the lateral frontal pole (FPl)—uniquely integrates information about the emotional unpleasantness and controllability of a future event, and that the strength of this integrated signal predicts better behavioral performance. These findings extend models of LPFC function to naturalistic, emotional contexts, and highlight the FPl as a key node for translating emotional and behavioral-control information into adaptive action.

## Introduction

> “Reason alone … can never oppose passion in the direction of the will” (Hume, 1739).

By characterizing goal pursuit as a process deeply intertwined with emotional experiences, Hume foreshadowed the need to study their integration. Emotional states inform the selection of goals and actions in everyday life and can sharpen or interfere with cognitive control depending on emotion-goal congruency (Braver et al., 2014; Chiew & Braver, 2011; Cohen et al., 2016; Gray & Braver, 2002). Successful goal-directed behavior and cognitive control are known to be supported by function of the lateral prefrontal cortex (LPFC) and its interconnected networks (Cole et al., 2016; Menon & D’Esposito, 2022; Waskom et al., 2014). Recent work underscores that emotional valence sculpts the representation of action goals in anterior LPFC (Lapate et al., 2022), and conversely, that LPFC function governs the influence of emotional stimuli on motor behavior (Bramson et al., 2020; Lapate et al., 2024). However, whether these recently uncovered properties of LPFC organization for behavioral control during emotional processing also extend to the anticipation of future emotionally-relevant events remains unknown. Here, we investigated the representational properties and temporal dynamics of affect and action-control coding in distinct LPFC regions during a powerful, naturalistic emotion induction—a threat of shock paradigm.

Prominent theories of the organization of control signals in LPFC postulate a rostro-caudal axis of abstraction along this region (Badre & D’Esposito, 2007; Bunge et al., 2005; Christoff & Gabrieli, 2000; Koechlin et al., 2003). Specifically, the lateral frontal pole (FPl) is thought to provide control signals that support temporally extended, abstract goal representations (Badre & Nee, 2018). Accordingly, FPl often shows ramping activation during prolonged, sequential control tasks (Desrochers et al., 2015, 2019). When temporal control is required, FPl signals have been hypothesized to inform mid-LPFC representations of context-sensitive action goals and rules (Nee & D’Esposito, 2017), which in turn guide action preparation and execution in caudal LPFC (e.g., premotor cortex (PMd)) (Binkofski & Buccino, 2006; Nakayama et al., 2008). The flexible integration of action control signals with time- and goal-relevant information is thought to promote adaptive responding across changing contexts (Badre et al., 2021).

Supporting a privileged role for FPl in goal-directed action in emotional contexts, recent anatomical evidence suggests that this region preferentially receives amygdala projections relative to other LPFC regions via the ventral amygdalofugal pathway (Folloni et al., 2019; Ghashghaei et al., 2007; Medalla & Barbas, 2010). Stronger amygdalofugal–FPl connectivity has been associated with increased emotion-driven influence on action control (Bramson et al., 2020), suggesting that FPl may be ideally located to integrate emotional information and LPFC-dependent action goals, which agrees with microstructural evidence identifying the frontopolar cortex as one of the most integrative regions within PFC (Badre & D’Esposito, 2009; Badre & Nee, 2018; Jacobs et al., 2001). Evidence for such integration was reported by Lapate and colleagues (2022), who identified significantly stronger conjunctive representations of emotional valence and action goals in FPl than in other LPFC regions during an emotion-dependent cognitive control task (Go/No-Go) (Lapate et al., 2022). Moreover, the strength of conjunctive emotion-action-goal representations in FPl correlated with better task performance (Lapate et al., 2022), aligning with extant theoretical and empirical work suggesting that higher-order, conjunctive representations may facilitate flexible context-sensitive behavior (Badre et al., 2021; Fusi et al., 2016; Kikumoto et al., 2024; Kikumoto & Mayr, 2020). Collectively, these studies suggest that FPl may be uniquely positioned to integrate amygdala-originated emotion signals with ongoing action goals to modulate cognitive control signals in LPFC.

While compelling, extant evidence for a privileged role of FPl function in emotion–action goal integration has largely come from studies employing relatively mild visual emotional stimuli (e.g., facial expressions), which also served as the imperative stimuli for action (i.e., they were action cues themselves) in those studies. Therefore, prior work leaves unclear whether the above-described affective coding properties in FPl extend beyond emotional-stimulus viewing to the anticipation of imminent, future emotionally potent events. Moreover, whether affective coding in FPl differs as a function of controllability over aversive events remains unknown. To that end, the present study examined the dynamic recruitment and representational properties—including the integration of affective and goal-relevant factors—in LPFC using multivariate analyses of fMRI data obtained during a naturalistic, self-relevant emotion induction procedure requiring goal-directed action: a threat-of-shock task.

Here, participants tracked a prolonged (18 s) countdown to possible shock administration. Shock unpleasantness and controllability were orthogonally manipulated, whereby a visual cue indicated whether a time-sensitive, precise motor response could prevent either a mild or unpleasant shock. In light of previous work uncovering functional and regional specificity in emotional-information coding in FPl, we hypothesized that FPl would show greater engagement during unpleasant (*vs.* mild) threat anticipation, and that FPl threat coding would be significantly stronger than in caudal LPFC regions (mid-LPFC and PMd). We also examined whether threat unpleasantness modulated the often-observed ramping in FPl during sequential events (Desrochers et al., 2015, 2019). Based on our previous findings (Lapate et al., 2022), we predicted that conjunctive representations of threat unpleasantness and controllability would be found in FPl and be stronger in this region than in caudal LPFC. Moreover, we predicted that stronger conjunctive representations of threat unpleasantness and controllability in FPl would be associated with better task performance (Kikumoto & Mayr, 2020; Lapate et al., 2022). Lastly, given recent work elucidating a pathway from the amygdala to FPl (Bramson et al., 2020; Folloni et al., 2019), we hypothesized that the strength of amygdala–FPl functional coupling would be associated with conjunctive representations of threat unpleasantness and controllability in FPl.

## Materials and Methods

### Participants

Sixty-seven participants were recruited at the University of California, Santa Barbara (UC Santa Barbara) (*M_age_* = 20.59 years; *SD_age_* = 2.00; range = 18 – 34 years; 51 female). The sample size was chosen based on the sample size used in a recent investigation of regional specificity in LPFC function during emotion-based cognitive control, which used representational similarity methods like the ones adopted in the current study) (Lapate et al., 2022). In that study, sample size was n = 37. We targeted a higher sample size (minimum n = 60) with the goal of maximizing power to replicate these effects in accordance with recent recommendations of n = 60 as the sample size required to detect a null effect with repeated-measures analysis (Brysbaert, 2019). All participants were healthy, with normal or corrected-to-normal vision, and no self-reported history of psychiatric or neurological disorders. Written informed consent was obtained from every participant. All study procedures were approved by the UC Santa Barbara Human Subjects Committee. Participants were compensated monetarily for their participation.

### Procedure

#### Overview

Following informed consent and MRI safety screening, physiological sensors were applied to participants and electrical shock calibration was conducted. The Active Escape task was completed during ~60 minutes of fMRI data collection. Upon completion, participants were paid for their time.

#### Experimental design and statistical analysis

##### Active Escape Task

In the MRI scanner, participants completed the Active Escape Task (adapted from Hur et al., 2020) which consisted of seven functional runs (lasting ~8 minutes each). Each functional run contained twelve trials, equally divided between “Controllable Unpleasant”, “Uncontrollable Unpleasant”, “Controllable Mild”, and “Uncontrollable Mild” trials. Trials were pseudorandomly presented, ensuring no consecutive repetitions of the same trial type **(Figure 1A)**. Each trial began with a fixation cross displayed for 500ms, followed by a 1000ms presentation of a cue letter and background color indicating threat unpleasantness and controllability. A cue letter “O” denoted a Controllable trial, whereas “X” denoted an Uncontrollable trial. Threat unpleasantness was denoted by the background color: red for Unpleasant shocks and blue for Mild shocks. Following the cue screen, an 18-second countdown began. The countdown period included the presentation of descending numbers (18, 17, 16…) flashed every second. The trial type (e.g., ‘Controllable Unpleasant’) was displayed as small text on the right side of the screen during the countdown. Upon completion of the countdown, participants were prompted to make a motor response using a handheld joystick to move a center cursor toward a target circle, which was randomly positioned in one of the four corners of the screen. Participants had 940ms to complete the response. The duration of the motor response window was determined following extensive in-scanner behavioral piloting, which indicated that 940ms reliably yielded accuracy rates between 60 - 80%. Successful motor responses (i.e., reaching the target circle within the allotted time) prevented shock administration during Controllable trials. A shock was delivered regardless of performance in Uncontrollable trials. Participants were instructed to respond on every trial, and were asked to make their best effort reaching the target circle irrespective of whether the trial was Controllable or Uncontrollable. Following conditional shock administration, a fixation cross was presented for 2000ms, followed by two questions assessing participants’ emotional experience. First, participants were asked to rate the intensity of their emotional experience during the countdown epoch: “*How intense was your emotional experience in this trial?”* on a scale of *1 (Not at all)* to *4 (Extremely)*. Next, they reported their confidence on their self-reported emotional evaluations: *“How certain are you in your response?”* on a scale of *1 (Not at all certain)* to *4 (Completely certain)* (data not reported here). For both questions, participants had up to 5000ms to respond. A variable 4000–6000ms intertrial interval followed. The Active Escape task totaled 84 trials divided into 7 blocks and took ~60 minutes to complete.

**Figure 1.**
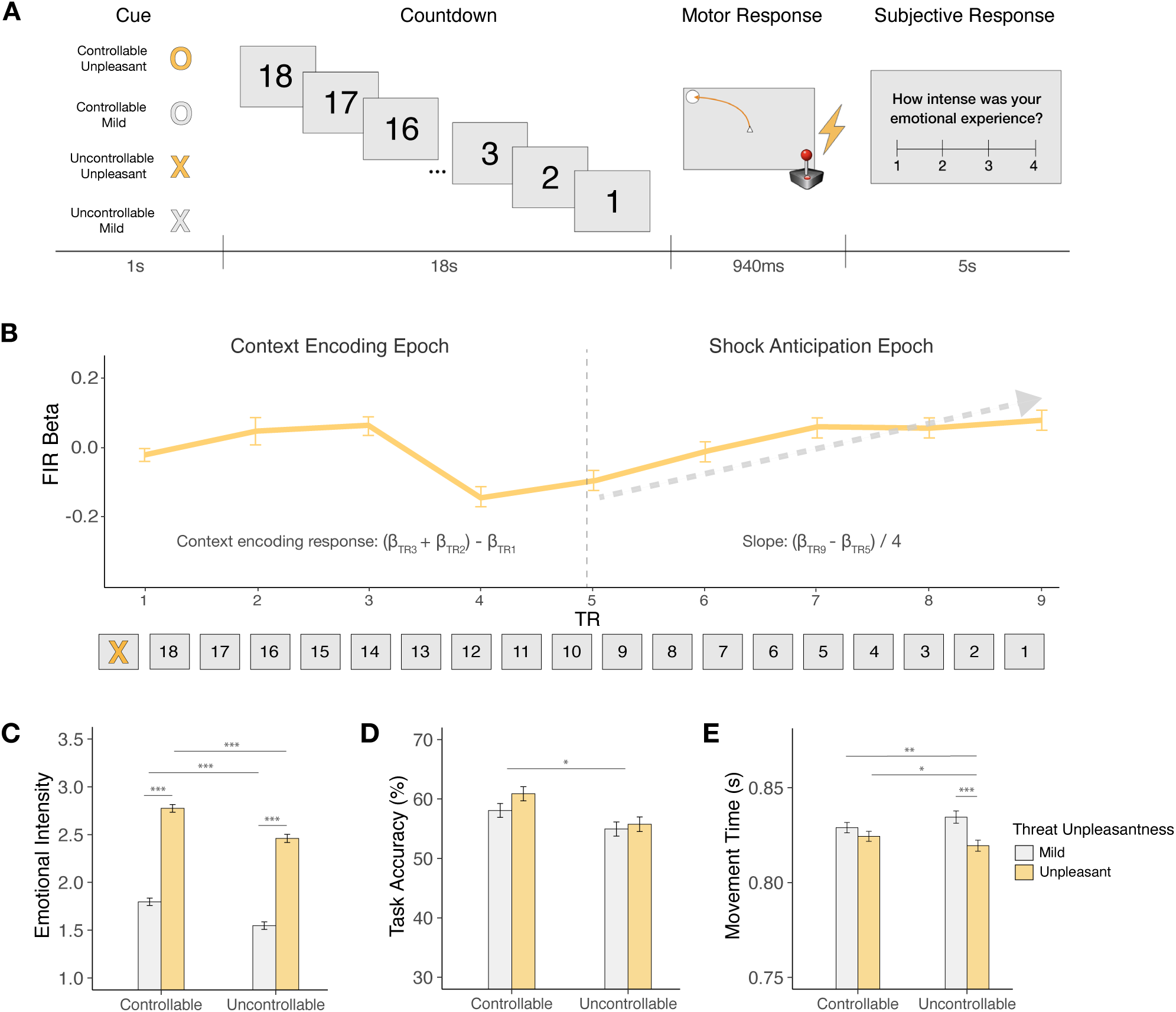
Design and validation of the Active Escape task. (**A**) *The Active Escape task*. In this threat-of-shock paradigm, participants tracked a prolonged countdown to Mild or Unpleasant shock administration, which was manipulated orthogonally in relation to threat controllability (Controllable *vs*. Uncontrollable). In Controllable trials, shocks could be avoided by making a successful motor response, whereas shocks were administered regardless of performance in Uncontrollable trials. At the start of each trial, participants saw a cue that informed them of the trial type (Controllable-Unpleasant, Controllable-Mild, Uncontrollable-Unpleasant, or Uncontrollable-Mild). Following each trial, participants reported the intensity of their emotional experience during the countdown. (**B**) *Neural Dynamics of Threat Anticipation.* FIR basis functions were used to estimate the magnitude of BOLD responses during the 18s countdown. The averaged FIR trace across FPl, mid-LPFC, and PMD is shown (collapsed across task conditions). Early and late neural dynamics were summarized over the first half (“context encoding epoch”) and the second half (“threat anticipation epoch”) of the countdown, referred to as a context encoding response metric, and a slope metric, respectively (*see Methods*). (**C**) *Emotional Intensity Ratings.* Participants reported experiencing greater emotional intensity during Unpleasant threat anticipation, particularly when anticipating Controllable (*vs*. Uncontrollable) Unpleasant threat. (**D**) *Task Performance.* Response accuracy was higher in Controllable, relative to Uncontrollable trials. (**E**) *Movement Time.* Participants were quicker to make successful motor responses when anticipating Unpleasant (*vs*. Mild) threat, particularly following Uncontrollable trials. Error bars denote the within-subjects 95% confidence interval for each condition, computed using Morey’s (2008) method (Morey, 2008). \**p* < 0.05, \*\**p* < 0.01, \*\*\**p* < 0.001.

##### Shock Stimuli

Electrical shocks were calibrated to each participant’s personal tolerance level (procedures adapted from Hur et al., 2020). Initial calibration was performed with participants lying in the scanner bed, but outside of the magnet bore. Electrical stimuli were generated using an MRI-compatible constant-voltage stimulator system (STM100C; Biopac Systems). Stimuli were delivered using MRI-compatible electrodes (EL509; Biopac Systems) placed on the inside of participants’ right wrist. Mild calibration: Participants received a low-intensity shock (2.5 Volts) and were asked whether the shock was “reliably detectable” and whether it was “at all unpleasant” (Hur et al., 2020). If participants reported the shock was not reliably detectable, the intensity was increased by 0.5 V, and the questions were repeated. Conversely, if the shock was reported as detectable and unpleasant, the intensity was decreased by 0.5 V, and the process was repeated. This iterative adjustment continued until participants reported the shock as reliably detectable but not at all unpleasant. The resulting stimulation intensity served as the Mild shock during the Active Escape task (M = 21.24 V, SD = 4.71 V, range = 10 - 35 V). Unpleasant shock calibration: Participants received a 26.25 V shock and were asked whether the shock was as “unpleasant as they were willing to tolerate” (Hur et al., 2020). If participants said no, the shock intensity increased by 2.5 V and the question was repeated. This process continued until participants reported that the shock was as unpleasant as they were willing to tolerate. This level of electrical stimulation served as the Unpleasant shock in the Active Escape task (M = 44.60 V, SD = 17.42 V, range = 26.25 – 103.75 V). A follow-up calibration was performed inside of the magnet bore, in which participants began with the shock intensity obtained in their initial calibration to confirm the Mild and Unpleasant shock levels previously obtained were still adequate.

##### Active Escape metrics and behavioral analyses

Task accuracy was calculated as the proportion of successful motor responses across trials. Movement time was measured as the time taken to successfully reach the target circle (correct trials only). We examined whether subjective emotional intensity, task accuracy, and movement time were modulated by threat unpleasantness and controllability using mixed effects models (with participant and run entered as random factors and threat unpleasantness and controllability entered as random slopes) in R (lme4 package (Bates et al., 2015), as well as *anova* (*stats* package (R Core Team (2013)) and *emmeans* (https://github.com/rvlenth/emmeans) packages for follow-up contrasts.

### Functional MRI Methods

#### Image acquisition

Neuroimaging data were acquired in the UC Santa Barbara Brain Imaging Center with a Siemens Prisma 3T MRI scanner equipped with a 64-channel radio frequency head coil. Whole-brain Blood Oxygenation Level-Dependent (BOLD) fMRI data were obtained using a T2*-weighted 2 accelerated multiband echoplanar imaging sequence (54 axial slices, 2.5mm^3^ isotropic voxels; 80×80 matrix; TR = 1900ms; TE = 30ms; flip angle = 65°; 250 image volumes/run). High-resolution anatomical scans were acquired using a T1-weighted magnetization-prepared rapid gradient echo (MPRAGE) sequence at the beginning of the session for spatial normalization (TR = 2500ms; TE = 2.22ms; flip angle = 7°), followed by a gradient echo field map (TR = 758ms; TE1 = 4.92ms; TE2 = 7.38ms; flip angle = 60°).

#### fMRI data preprocessing

Functional neuroimaging data were preprocessed using FEAT (FMRI Expert Analysis Tool) Version 6.00, implemented in FMRIB’s Software Library (Jenkinson et al., 2012; Smith et al., 2004). Preprocessing steps included applying high-pass temporal filtering with a 100s cutoff, inclusion of standard and extended motion correction regressors obtained using MCFLIRT, spatial smoothing using a Gaussian kernel of 5mm, slice-time correction, and adding regressors of non-interest created from a matrix of points representing displacement of 0.5 mm or greater to control for movement-confounded activation. Brain extraction was carried out through Advanced Normalization Tools (ANTs) skull-stripping algorithm (Avants et al., 2009). Functional images were co-registered to participant’s T1-weighted anatomical image using a linear rigid body (6-DOF) transform while maintaining native functional resolution (2.5 mm^3^ isotropic).

#### Regions of Interest

##### Subcortical

The amygdala was defined using the Harvard Oxford atlas mask (Jenkinson et al., 2012; Smith et al., 2004) thresholded at 50% and registered from MNI to participants’ structural space using FNIRT (10 mm warp) while maintaining native resolution (2.5 mm^3^ isotropic).

##### Cortical

Prefrontal ROIs [FPl, mid-LPFC (9-46d and 9-46v), and PMd] were obtained from the Oxford PFC Consensus Atlas (http://lennartverhagen. com/; Neubert et al., 2014; Sallet et al., 2013), thresholded at 25% and were then registered to participants’ individual anatomical scans using Freesurfer (Reuter et al., 2012). Vertex coordinates within each ROI were transformed into native space, and volumetric ROI masks were created by projecting half of the cortical thickness at each vertex. A functional voxel was included if at least 50% of its volume was covered by the label, with intersected voxels being labeled accordingly (Lapate et al., 2022). We selected FPl, mid-LPFC, and PMd ROIs as key nodes along the LPFC rostrocaudal axis. mid-LPFC was defined as BA 9-46d and BA 9-46v, based on prior research identifying this region as a critical node for the integration of cognitive control information (Lapate et al., 2022; Lapate et al., 2024; Miller et al., 2022).

#### Finite Impulse Response

To examine the temporal dynamics of distinct LPFC regions during threat anticipation, we used Finite Impulse Response (FIR) basis functions, which provide a model-free estimation of BOLD signals over time (Friston, 2005; Grupe et al., 2018). In FEAT (Jenkinson et al., 2012; Smith et al., 2004), we modeled the time series data using 10 basis functions for a 19s window, corresponding to each TR during the countdown. We then extracted beta activation estimates for each individual, TR, and for each level of threat unpleasantness and controllability, and used mixed-effects models (with subject modeled as a random factor and unpleasantness and controllability as random slopes, as described above) to determine BOLD modulation as a function of region, TR, threat and controllability.

To interrogate the dynamics of LPFC engagement as estimated by the FIR model, we compared the model fits of first, second, and third-order polynomials to determine the best-fitting model of the 18s countdown time course. This model comparison indicated that third-order polynomials best captured the shape of BOLD activation in FPl, mid-LPFC, and PMd during the countdown period. This informed the division of the countdown into two epochs (henceforth called “context encoding” and “shock anticipation” epochs). We created two metrics to capture these distinct BOLD responses in the first and second halves of the countdown: (1) The *context encoding response* captured neural activation at the onset of the trial, upon the presentation of the trial cue that conveyed the task condition (i.e., context) that the participants were in (i.e., the threat unpleasantness and controllability of the present trial). This metric was computed by identifying and summing the activation peaks for FPl, mid-LPFC, and PMd, which occurred at TR_2_ and TR_3_, and subtracting the baseline activation at TR_1_ [(TR_2_ + TR_3_) − TR_1_]. (2) The final *shock anticipation* metric was computed by taking by the slope of FIR betas from TRs 5 to 9 [Δy / Δx : (TR_9_ − TR_5_) / 4]. TR 5 was identified as the trough in activation for FPl, mid-LPFC, and PMd, and TR 9 was the final TR of the countdown before participants would be prompted to make the motor response. This metric was computed to capture the ramping dynamics occurring in the last half of the countdown epoch, as the electric shock (and future motor response) approached (Desrochers et al., 2015, 2019) (see **Figure 1B** for LPFC beta coefficients plotted by TR, collapsed across task conditions).

#### Representational similarity analysis

We used representational similarity analysis (RSA) (Diedrichsen & Kriegeskorte, 2017; Kriegeskorte et al., 2008) to examine the structure of threat unpleasantness and controllability representations along the rostro-caudal axis of the LPFC. Following our previous work (Lapate et al., 2022), we obtained voxel-wise and trial-wise BOLD activation parameters estimates using the Least-Squares All general linear model (GLM) approach (Mumford et al., 2012) and FEAT modeling in FSL (Jenkinson et al., 2012; Smith et al., 2004). Single trials were modeled using a double-gamma hemodynamic response function. Parameter estimates extracted from each ROI were regularized with multivariate noise normalization by obtaining an estimate of the noise covariance from the residuals of the GLMs from each ROI. This matrix was then regularized using the optimal shrinkage parameter, inverted, and multiplied by the vector of betas for each trial (Ballard et al., 2019; Walther et al., 2016). This approach removes spurious correlations between voxels caused physiological and environmental noise (Walther et al., 2016).

Factorial RSA uses template matrices to test whether the intertrial similarity structure of multivoxel activity patterns is explained by task factors independently (i.e., threat unpleasantness and/or controllability) or by their interaction (i.e., threat unpleasantness * controllability). To construct the neural similarity matrix for each ROI, we used a cross-validation approach by computing Pearson correlation coefficients between multivoxel patterns for all possible trial pairs across runs, excluding trials from within the same run (Lapate et al., 2022). This resulted in one trial-wise neural similarity matrix per participant and ROI. Then, we fit simultaneous multiple regression models with condition-specific template matrices to examine whether the similarity of multivoxel patterns from each ROI’s neural similarity matrix was modulated by threat unpleasantness, controllability—and/or their interaction, consistent with higher-order, conjunctive representations (Badre et al., 2021; Kikumoto & Mayr, 2020). Our primary set of model matrices tested for *discriminative* structures across each pairing of threat unpleasantness and controllability levels. In other words, the discriminative model matrices tested whether the representational distance in a high-dimensional space was differentially influenced by threat unpleasantness or controllability (for instance, whether the multivariate distance between neural activity patterns for a given ROI differed between Unpleasant and Mild shock trials, or between Controllable and Uncontrollable trials). Critically, the interactive discriminative matrix of threat unpleasantness and controllability (threat unpleasantness * controllability) tests whether the representational distance of the levels of a given task factor (e.g., threat unpleasantness) is modulated by another (e.g., controllability)—for instance, whether Unpleasant shock trials are represented differently from Mild shock trials depending on controllability (Ballard et al., 2019; Diedrichsen & Kriegeskorte, 2017; Kriegeskorte et al., 2008; Lapate et al., 2022).

We also included traditional similarity template matrices in our regression model, which tested for intertrial similarity while assuming equal representational distance across conditions (e.g., equivalent pattern distances across Unpleasant versus Mild shock trials). We fit these five template matrices on the neural similarity matrices using a multiple regression mixed-model framework, with subject modeled as a random factor, and each unique template matrix included in the subject error term (per epoch). This allowed us to test whether FPl, mid-LPFC, and PMd represented the interaction of threat unpleasantness or controllability (i.e., conjunctive representations), *vs.* whether these regions represented threat unpleasantness and/or controllability dimensions (Unpleasant *vs.* Mild; Controllable *vs.* Uncontrollable) independently (Lapate et al., 2022). As we found significant conjunctive representations in FPl, we correlated FPl RSA betas with task accuracy across subjects to examine whether the strength of conjunctive representations was associated with behavioral performance (Lapate et al., 2022). Because outliers were observed in the RSA betas, we used Spearman’s ρ to conduct correlational analyses. When pertinent, differences in strength of correlation coefficients were tested using CoCor (Diedenhofen & Musch, 2015).

#### Cross-condition generalization performance analysis

To characterize the generalizability of LPFC representations, we next conducted cross-condition generalization performance (CCGP) analysis (Bernardi et al., 2020). To that end, using multivariate pattern analysis (MVPA) (with multivariate-noise normalized single-trial betas, as described above), we tested whether a Logistic Classifier trained on a task condition set *within* a particular experimental factor level (e.g. Unpleasant *vs*. Mild shock conditions during Controllable trials) could generalize to untrained levels of another experimental factor (e.g., Unpleasant *vs*. Mild shock during Uncontrollable trials). Specifically, for each subject and ROI, we used a multivariate logistic regression model (l2 penalty; C = 1) to iteratively train the classifier and assessed classifier performance using a leave-one-run-out cross-validation scheme. Classification performance was evaluated using the area under the curve (AUC; i.e., where 0.5 is chance performance). This was implemented in Nilearn (Abraham et al., 2014) using the parameter (class_weight = ‘balanced’) to automatically adjust weights according to class frequencies in the input data. To determine whether classification accuracy differed from chance, we combined run-wise classifier AUCs (– 0.5) across subjects using a mixed-model approach, where subject and run were entered as random factors, and tested whether the intercept differed significantly from 0.

This was repeated to account for every possible combination of split training and testing sets, using a leave-one-run-out cross-validation scheme. We obtained CCGP for all balanced combinations of threat unpleasantness and controllability (per epoch). CCGP is quantified as the averaged cross-validated performance across levels, where above-chance classifier performance indicates coding generalization across conditions (Bernardi et al., 2020). In general, higher CCGP is thought to reflect greater generalization or coding abstraction, and be related to lower-dimensional representations (Bernardi et al., 2020; Bhandari et al., 2024).

#### Functional connectivity analysis

To test whether LPFC task representations were modulated by emotion-related amygdala signals, we used psychophysiological interaction (PPI) analysis (O’Reilly et al., 2012). To that end, we extracted the mean time series data for the amygdala, which was used as the seed region. We then ran a FEAT first-level model analysis that included paired threat unpleasantness and controllability regressors, the demeaned amygdala timecourse, as well as the interaction between this timecourse and the task condition regressors. The resulting parameter estimates from this interaction regressor, indexing task-dependent functional connectivity of the amygdala with each voxel of the brain, were extracted for each subject, functional run, and ROI.

To test whether there was significant coupling between the amygdala and FPl during the Active Escape task, we conducted a mixed-effects model on the resulting PPI betas with threat unpleasantness and controllability as fixed factors (including subject and run as random factors). Thus, this model tested whether the intercept differed significantly from zero, as well as whether amygdala functional connectivity significantly differed as a function of task condition. Following a significant association between amygdala–FPl PPI betas and FPl conjunctive RSA betas, we additionally correlated PPI and RSA betas across-subjects to test the regional specificity of this association. When relevant (e.g., when comparing whether the magnitude of an association differed across distinct LPFC regions), we tested for the difference in correlation coefficients using CoCor (Diedenhofen & Musch, 2015).

## Results

### Controllable escape from threat modulates experienced emotion and facilitates task performance

We first verified that our threat-of-shock paradigm produced changes in subjective emotion and goal-dependent task performance. As predicted, threat unpleasantness robustly increased self-reported emotional intensity (*F* = 283.30, *p* < 0.001), such that participants reported significantly greater emotional intensity when anticipating Unpleasant relative to Mild shocks (*B* = 0.94 (SE = 0.06), *t* = 16.83, *p* < 0.001, *d_z_* = 2.02, 95% CI [0.83, 1.05]). A main effect of controllability was also observed (*F* = 35.47, *p* < 0.001), with higher emotional intensity ratings for Controllable relative to Uncontrollable trials (*B* = 0.28 (SE = 0.05), *t* = 5.96, *p* < 0.001, *d_z_* = 0.71, 95% CI [0.19, 0.38]). Importantly, these effects were qualified by a two-way Threat Unpleasantness * Controllability interaction (*F* = 4.46, *p* = 0.04), wherein Controllable Unpleasant shock trials elicited higher emotional intensity than all other trial types (*Controllable Unpleasant vs.* Uncontrollable Unpleasant: *B* = 0.32 (SE = 0.05), *t* = 6.32, *p* < 0.001, *d_z_* = 0.66, 95% CI [0.19, 0.45]; *vs.* Controllable Mild: *B* = 0.98 (SE = 0.06), *t* = 16.75, *p* < 0.001, *d_z_* = 1.98, 95% CI [0.83, 1.13]; *vs.* Uncontrollable Mild: *B* = 1.23 (SE = 0.07), *t* = 16.81, *p* < 0.001, *d_z_* = 2.03, 95% CI [1.04, 1.42]) (**Figure 1C**). When examining the impact of threat unpleasantness and controllability on task accuracy, we found that controllability benefitted performance (*F* = 6.61, *p* = 0.01), with participants performing more accurately in Controllable than Uncontrollable trials (*B* = 0.04 (SE = 0.02), *t* = 2.57, *p* = 0.01, *d_z_* = 0.32, 95% CI [0.01, 0.07]). Other effects did not reach significance [threat unpleasantness main effect: *p* = 0.15; Threat Unpleasantness * Controllability interaction: *p* > 0.49] (**Figure 1D**). Of note, Unpleasant (*vs.* Mild) threat anticipation led to faster movement times (*F* = 15.40, *p* < 0.001; *B* = −0.02 (SE = 0.003), *t* = −3.925, *d_z_* = −0.45, 95% CI [−0.02, −0.01]), particularly during Uncontrollable trials (Threat Unpleasantness * Controllability interaction: *F* = 8.26, *p* = 0.004; Uncontrollable Unpleasant *vs.* Mild: *B* = −0.02 (SE = 0.004), *t* = −4.78, *p* < 0.001, *d_z_* = −0.43, 95% CI [−0.03, −0.01]; Uncontrollable Unpleasant *vs.* Controllable Unpleasant: *B* = − 0.009 (SE = 0.003), *t* = −2.81, *p* = 0.03, *d_z_* = −0.13, 95% CI [−0.02, −0.001]) **(Figure 1E)**.

In summary, anticipating unpleasant controllable shocks evoked greater emotional intensity than anticipating either mild or uncontrollable shocks. Movement times were faster following the anticipation of unpleasant (*vs*. mild) shocks, and controllable (*vs.* uncontrollable) threats enhanced task accuracy. Taken together, these results indicate that both the anticipation of threat and its controllability increased the intensity of subjective emotional experiences, and benefitted task performance.

Next, we examined the temporal dynamics of threat-of-shock responding in distinct LPFC regions along the rostro-caudal axis to test whether FPl showed stronger coding of threat unpleasantness relative to caudal regions. To that end, we fit FIR basis functions over the 18s task countdown, and extracted early and late BOLD responses during threat anticipation as indexed by (1) the overall magnitude of the initial encoding of trial condition (FIR amplitude during the ‘context encoding’ epoch), and (2) the ramping of the BOLD response in the epoch preceding potential shock administration (FIR slope in the ‘shock anticipation’ epoch; for details, see *Methods*).

### FPl shows pronounced emotion-modulated responding during initial threat coding

During the context encoding epoch, threat anticipation robustly increased activation across LPFC as evidenced by a significant main effect of threat unpleasantness, with greater activation during the anticipation of Unpleasant *versus* Mild shocks (*F* = 24.61, *p* < 0.001). Critically, the magnitude of threat-related coding significantly differed across regions along the LPFC rostro-caudal axis [Threat Unpleasantness * Region interaction: *F* = 5.36, *p* = 0.005; other interactions were n.s., *p*s > 0.18] (see **Figure 2A** for threat responding (Δ Unpleasant − Mild) plotted by region). This two-way interaction reflected that the finding that FPl showed the strongest modulation by threat unpleasantness (*B* = 0.34 (SE = 0.06), *t* = 5.77, *p* < 0.001, *d_z_* = 0.62, 95% CI [0.22, 0.47]), followed by mid-LPFC (*B* = 0.19 (SE = 0.06), *t* = 3.25, *p* = 0.001, *d_z_* = 0.57, 95% CI [0.12, 0.27]), and PMd (*B* = 0.12 (SE = 0.06), *t* = 2.06, *p* = 0.04, *d_z_* = 0.31, 95% CI [0.05, 0.19]), respectively. Of note, the magnitude of LPFC engagement during the context encoding epoch was associated with stronger self-reported emotional intensity in FPl (*⍴* = 0.26, *p* = 0.04) and in mid-LPFC (*⍴* = 0.34 *p* = 0.01) (see **Figure S1**). In summary, LPFC was robustly engaged during the initial encoding of future threat, with FPl showing the greatest amplitude modulation by threat unpleasantness—an effect that progressively attenuated along the rostro-caudal axis.

**Figure 2.**
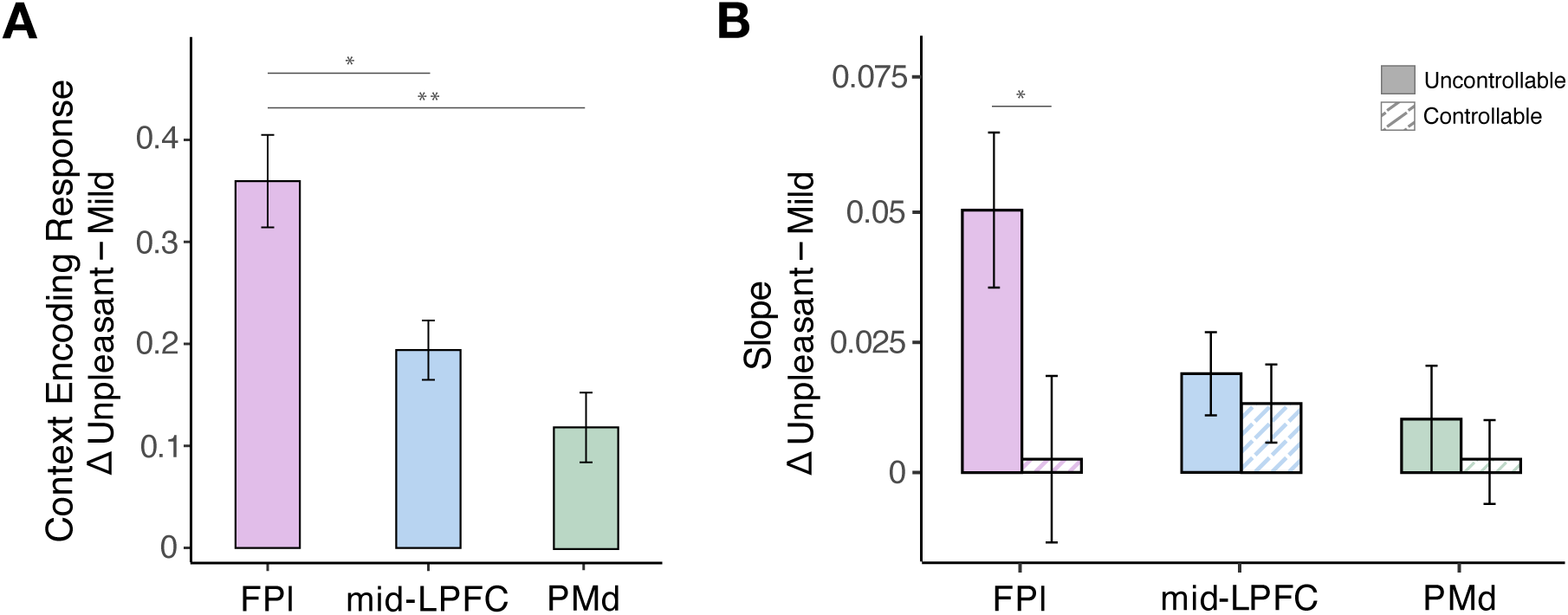
Regionally specific recruitment of LPFC during prolonged threat of shock anticipation and association with subjective experience. (**A**) During the initial context-encoding epoch, anticipation of Unpleasant (*vs.* Mild) threat elicited robust engagement across LPFC ROIs (FPl, mid-LPFC, and PMd); moreover, FPl showed stronger discrimination of Unpleasant (*vs.* Mild) threat relative to mid-LPFC and PMd. (**B**) During the later shock anticipation epoch, threat unpleasantness and controllability jointly modulated FPl function, as evidenced by robust ramping in FPl (*vs.* other LPFC regions) when anticipating Uncontrollable Unpleasant (*vs*. Uncontrollable Mild or Controllable) shocks. Error bars denote the 95% within-subject confidence interval standard error of the mean (Morey, 2008). \**p* < 0.05, \*\**p* < 0.01.

### FPl ramping during threat anticipation is modulated by threat unpleasantness and controllability

As FPl typically shows robust ramping during sequential processing and control tasks (Desrochers et al., 2015, 2019), we next tested whether FPl showed its characteristic activation ramping during the final countdown period preceding potential shock administration and tested whether this hypothesized ramping was sensitive to threat-relevant information (relative to other LPFC regions). To this end, we computed the slope of activation during the shock anticipation epoch (see *Methods*).

We found that the magnitude of ramping activation preceding potential shock administration significantly differed across LPFC regions (region main effect: *F* = 5.12, *p* = 0.006), such that the slope was largest in FPl (M = 0.05, SE = 0.011), followed by mid-LPFC (M = 0.04, SE = 0.005) and PMd (M = 0.03, SE = 0.007), respectively; FPl *vs.* PMd (*B* = 0.02, (SE = 0.006), *t* = 3.08, *p* = 0.006, *d_z_* = 0.17, 95% CI [0.004, 0.03]). Moreover, threat unpleasantness significantly modulated the magnitude of ramping across all three regions (threat unpleasantness main effect: *F* = 5.17, *p* = 0.03), with steeper slopes during the anticipation of an Unpleasant (*vs.* Mild) shock (*B* = 0.02 (SE = 0.007), *t* = 2.27, *p* = 0.02, *d_z_* = 0.29, 95% CI [0.002, 0.03]). This was further qualified by a significant Threat Unpleasantness * Controllability interaction (*F* = 4.34, *p* = 0.04), where the modulation of ramping by threat unpleasantness was stronger in Uncontrollable (*vs.* Controllable) trials (Uncontrollable Unpleasant − Mild: *B* = 0.03 (SE = 0.03), *t* = 3.06, *p* = 0.01, *d_z_* = 0.34, 95% CI [0.004, 0.05]), such that ramping was steepest in *Uncontrollable* Unpleasant trials. When examining these effects for each ROI, we found that Unpleasant (*vs*. Mild) shock trials produced steeper slopes in FPl [threat unpleasantness main effect: *F* = 4.08, *p* = 0.04] and mid-LPFC [threat unpleasantness main effect: *F* = 6.59, *p* = 0.01], but not PMd (*p* = 0.19). Likewise, the Threat Unpleasantness * Controllability interaction was found in FPl (Uncontrollable Unpleasant − Uncontrollable Mild: *B* = 0.05 (SE = 0.02), *t* = 2.88, *p* = 0.02, *d_z_* = 0.35, 95% CI [0.005, 0.09]), but not in mid-LPFC or PMd (all *ps* > 0.40) (although note that a Region * Threat Unpleasantness * Controllability interaction did not reach significance (*p* = 0.18)) (**Figure 2B**). (For additional analyses—including TR-by-TR and the modulation of activation by unpleasantness and controllability in other LPFC and mPFC ROIs, see **Figures S2** and **S3**). In summary, FPl showed pronounced ramping during unpleasant threat anticipation compared to other LPFC regions, which was steepest when the impending threat was uncontrollable.

### FPl holds conjunctive representations of threat unpleasantness and controllability

The above-reported findings—indicating that FPl function is modulated by both threat unpleasantness and controllability—raise the question of whether FPl codes for these distinct task features into integrated, putatively higher-order conjunctive representations (Badre et al., 2021; Ballard et al., 2019; Kikumoto & Mayr, 2020; Lapate et al., 2022). To address this, we conducted full-factorial representational similarity analysis (RSA) (Kriegeskorte et al., 2008). First, we computed neural similarity matrices for each ROI using trial-wise data (across runs). Next, we regressed two sets of template matrices on the neural similarity matrix to capture the distinct ways in which threat unpleasantness and controllability may be represented (cf. Ballard et al., 2019; Lapate et al., 2022). Our primary set of model matrices—referred to as condition *discrimination matrices*—tested whether representational distances differed between distinct levels of threat unpleasantness and controllability, as well as their interaction (i.e., Threat Unpleasantness * Controllability) (see **Figure 3A** for an illustration and *Methods* for further details). In particular, the interactive discrimination matrix tested whether representational distances between levels of one factor (e.g., Unpleasant *vs.* Mild trials) depended on levels of another factor (e.g., threat controllability). To account for potential inter-trial condition similarities—with equivalent representational distances across condition levels—we also included a set of *similarity matrices*. We fit these template matrices to the neural similarity matrix using a simultaneous multiple regression approach, which allowed us to determine whether the inter-trial similarity structure of multivoxel activity patterns in each ROI was best explained by threat unpleasantness and controllability independently, or by their interaction (cf. Lapate et al., 2022) (for details, see *Methods*).

**Figure 3.**
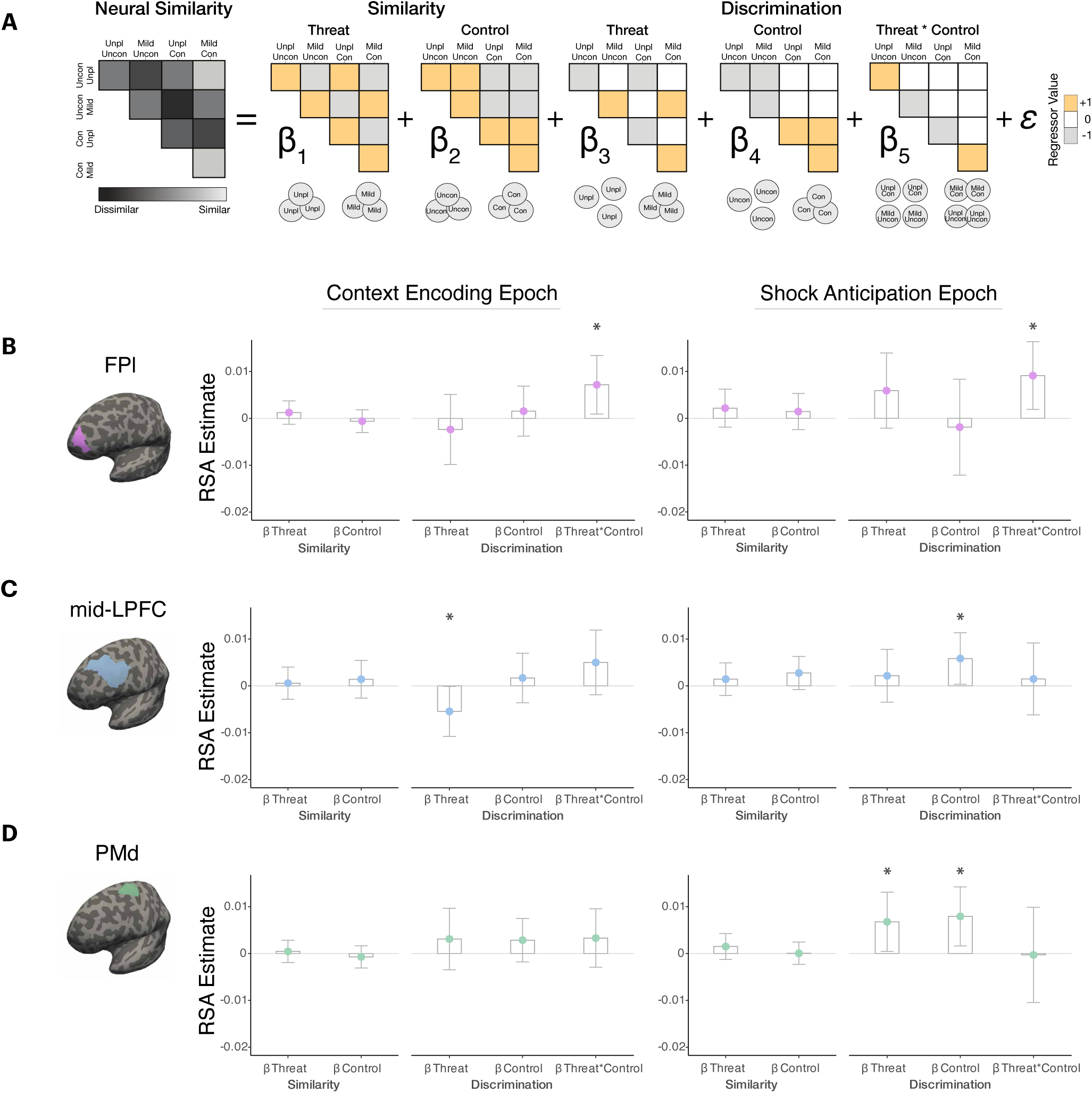
FPl holds conjunctive coding of threat-relevant information. (**A**) *RSA Matrices*. Condition-specific template matrices tested whether the correlation of multivoxel activity patterns across trials and across runs was best explained by threat unpleasantness, controllability, and/or their interaction (conjunctive threat * control representations). Template matrices modeled both equivalent representational distances across condition levels (similarity matrices) as well as differential representational distances across condition levels (discrimination matrices). Lower circles illustrate the representational structure captured by each regressor. (For brevity, threat unpleasantness and controllability factors are labeled in the abscissa as “threat” and “control”, respectively). (**B**) *FPl RSA Betas.* Representational similarity analyses revealed conjunctive representations of threat unpleasantness and controllability in FPl across both the context encoding and shock anticipation epochs. (**C**) *mid-LPFC RSA Betas.* Mid-LPFC held independent representations of threat unpleasantness during context encoding and controllability during shock anticipation. (**D**) *PMd RSA Betas.* PMd held independent representations of both threat unpleasantness and controllability during shock anticipation. \**p* < 0.05

Critically, we found evidence of conjunctive threat unpleasantness and controllability representations in FPl, as evidenced by the significant fit of the Threat Unpleasantness * Controllability discrimination matrix during both the context encoding epoch (*F* = 5.09, *p* = 0.03) and the shock anticipation epoch (*F* = 6.12, *p* = 0.02) (**Figure 3B**), consistent with the integration of threat unpleasantness and controllability information in this region. In contrast, mid-LPFC held independent representations of threat unpleasantness and controllability during the context encoding (*F* = 3.99, *p* = 0.048) and shock anticipation epochs (*F* = 4.36, *p* = 0.04), respectively (**Figure 3C**). Similarly, PMd held independent representations of threat unpleasantness (*F* = 4.42, *p* = 0.04) and controllability (*F* = 6.04, *p* = 0.02) as the threat became proximal–i.e., during the shock anticipation epoch (**Figure 3D**). Neither mid-LPFC nor PMd showed evidence of conjunctive emotion-control representations in either epoch [context encoding epoch: *p*s > 0.16; shock anticipation epoch: *p*s > 0.70].

Importantly, conjunctive threat unpleasantness and controllability representations in FPl were significantly stronger than in other LPFC regions during the shock anticipation epoch, thereby indicating regional specificity in the representation of integrated emotional and control-related information as threat nears in time (Region * Threat Unpleasantness * Controllability interaction: *F* = 3.22, *p* = 0.04; FPl *vs.* mid-LPFC: *B* = 0.008 (SE = 0.004), *t* = 1.89, *p* = 0.06, *d_z_* = 0.25, 95% CI [−0.0003, 0.02]; FPl *vs.* PMd: *B* = 0.01 (SE = 0.004), *t* = 2.41, *p* = 0.02, *d_z_* = 0.32, 95% CI [0.002, 0.02]). In summary, these findings uncover region-specific representational geometries in LPFC and suggest a privileged role for FPl in integrating unpleasantness and controllability during the anticipation of naturalistic, self-relevant threat.

Conjunctive representations are thought to be maintained in high-dimensional representational spaces, where the degree of task feature separation depends on the extent to which representations are context specific (Bernardi et al., 2020; Kikumoto et al., 2024; Rigotti et al., 2013). To characterize the generalizability of FPl conjunctive representations, we conducted a cross-condition generalization performance (CCGP) analysis (Bernardi et al., 2020), which quantifies how well a linear classifier trained to decode a task-relevant variable in one set of task conditions (e.g., specific level of a task factor) generalizes to a set of untrained conditions— thereby indexing the level of abstraction or context independence of the underlying representation (for details, see *Methods*). We found that while decoding of threat unpleasantness in FPl was above chance in the shock anticipation epoch (*M* = 54.16%, *B* = 0.004 (SE = 0.02), *t* = 2.03, *p* = 0.02, *d_z_* = 0.26, 95% CI [0.50, 0.58])—threat decoding did not generalize across controllability conditions in this region (*p*s > 0.16). This pattern is consistent with a high-dimensional structure in FPl, with threat unpleasantness and controllability represented in a context-specific manner as threat nears in time (Badre et al., 2021; Bernardi et al., 2020; Rigotti et al., 2013).

### FPl conjunctive representations correlate with task performance

By integrating multiple task features, conjunctive representations have been hypothesized to confer functional benefits, such as facilitating context-sensitive behavioral flexibility (e.g., Badre et al., 2021). We therefore examined whether conjunctive representations in FPl were associated with task performance—here, a time-sensitive, precise motor action that could prevent shock administration in Controllable trials. We found that stronger conjunctive coding of threat unpleasantness and controllability in FPl was associated with better task performance as Controllable (*vs.* Uncontrollable) Unpleasant shock trials neared in time (i.e., during the shock anticipation epoch) (*⍴* = 0.30, *p* = 0.03) (**Figure 4**). This association was not observed in mid-LPFC (*⍴* = −0.02, *p* = 0.87) or PMd (*⍴* = 0.05, *p* = 0.75) (see *Supplemental Materials* **Figure S4**). These findings suggest that FPl integration of threat unpleasantness and controllability may be functionally advantageous and guide behavior under emotionally salient contexts that require swift, goal-directed action.

**Figure 4.**
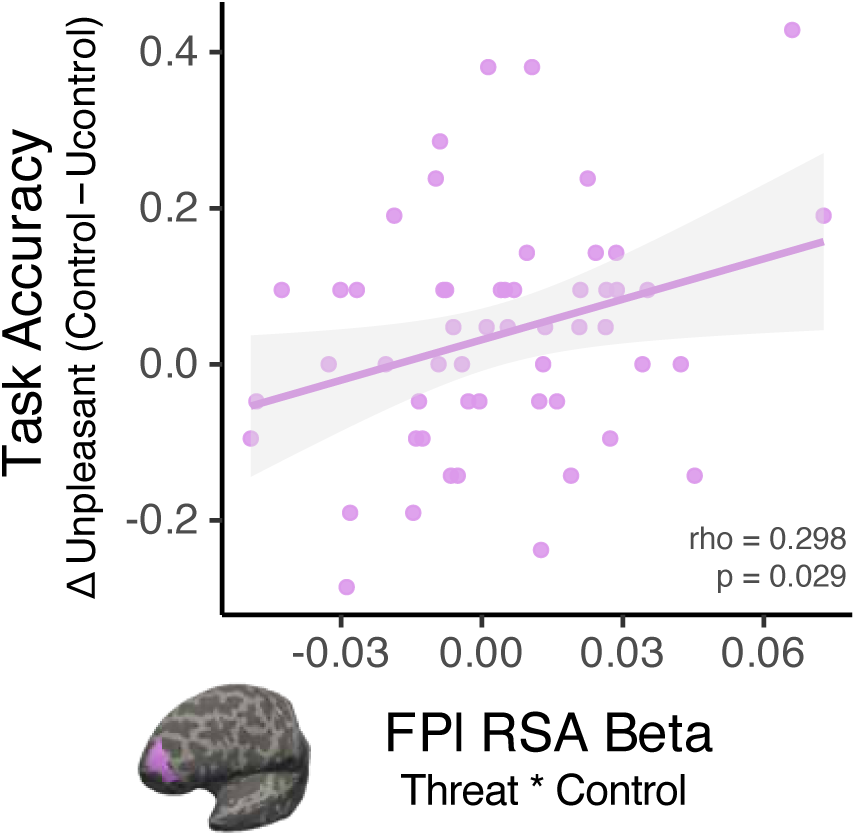
FPl conjunctive coding is associated with task accuracy. Stronger conjunctive representations of Threat Unpleasantness and Controllability in FPl correlated with higher task performance in Controllable Unpleasant (*vs.* Uncontrollable Unpleasant) trials as threat neared in time (during the shock anticipation epoch).

### Amygdala–FPl connectivity under threat may support integrated FPl representations

Recent anatomical evidence suggests that FPl has preferential access to amygdala projections relative to other LPFC regions (Bramson et al., 2020; Folloni et al., 2019; Ghashghaei et al., 2007; Medalla & Barbas, 2010). Thus, it is possible that conjunctive representations in FPl are in part informed by—and associated with—the strength of amygdala signaling. To examine this possibility, we performed a psychophysiological interaction (PPI) analysis with the amygdala as the seed region. This analysis revealed a significant modulation of amygdala–FPl coupling as a function of controllability, with more inverse amygdala–FPl coupling in Uncontrollable *vs*. Controllable trials (*F* = 4.88, *p* = 0.03) (**Figure 5A**).

**Figure 5.**
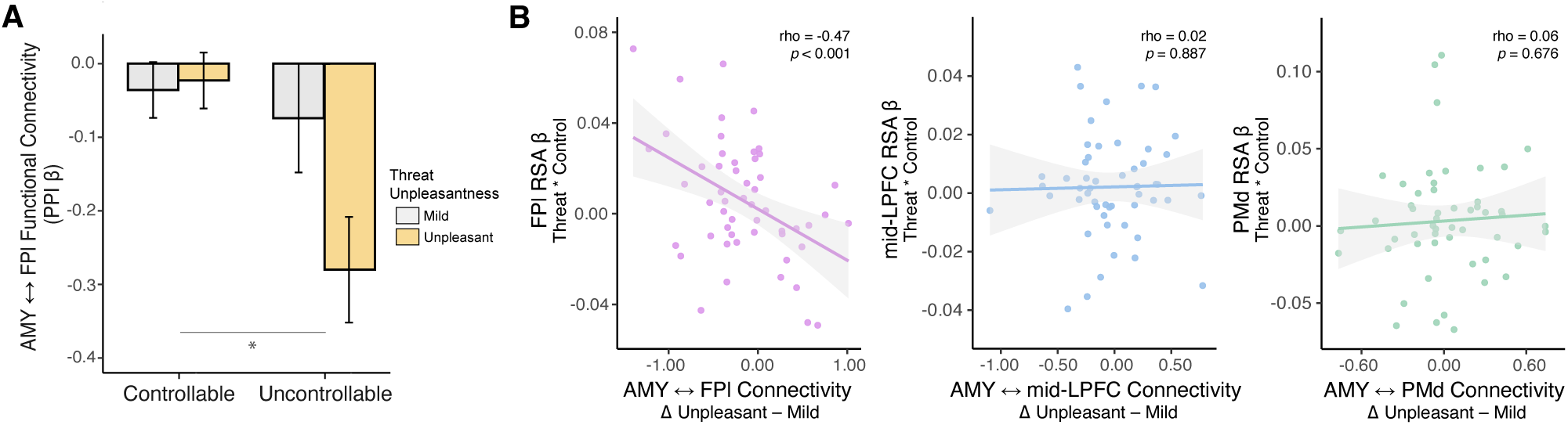
Amygdala–LPFC Functional Connectivity. (**A**) Controllability-modulated amygdala–FPl connectivity. A PPI analysis revealed that the magnitude of amygdala–FPl connectivity is modulated by controllability, with significantly more negative coupling during Uncontrollable (*vs*. Controllable) threat trials. (**B**) Associations between amygdala–LPFC functional connectivity and the strength of conjunctive RSA betas. Inverse amygdala– FPl coupling was associated with stronger conjunctive representations of threat unpleasantness and controllability in FPl (**left panel**). In contrast, the strength of amygdala coupling with mid-LPFC (**center panel**) or PMd (**right panel**) was not associated with the magnitude of conjunctive representations in those regions. Error bars denote the within-subjects 95% confidence interval for each condition, computed using Morey’s (2008) method (Morey, 2008).

Next, we examined whether amygdala–FPl coupling was associated with the strength of conjunctive representations in FPl. We found that stronger conjunctive coding in FPl was associated with more inverse amygdala–FPl coupling during the anticipation of Unpleasant (*vs*. Mild) shock trials (*⍴* = −0.47, *p* < 0.001). In contrast, amygdala coupling with other LPFC regions was not significantly modulated by threat unpleasantness (mid-LPFC *p* = 0.20; PMd *p* = 0.55) or controllability (mid-LPFC *p* = 0.06; PMd *p* = 0.07), Moreover, suggesting regional specificity of the association between amygdala coupling and the representational structure of threat unpleasantness and controllability to FPl, no significant associations were found between amygdala coupling and Threat Unpleasantness * Controllability representations in either mid-LPFC (*p* > 0.56) or PMd (*p* > 0.40) (**Figure 5B**). Importantly, the correlation between amygdala coupling and conjunctive unpleasantness-controllability coding was significantly stronger in FPl compared to its caudal counterparts (*r* difference FPl *vs.* mid-LPFC: *t* = 2.65, *p* = 0.01; FPl *vs*. PMd: *t* = 2.07, *p* = 0.04). Taken together, these results suggest that threat-modulated amygdala–FPl connectivity may support the formation and/or maintenance of conjunctive representations of threat unpleasantness and controllability in FPl.

## Discussion

By examining the temporal dynamics of LPFC engagement during the prolonged anticipation of naturalistic, self-relevant threat using univariate and multivariate fMRI analyses, we found that distinct LPFC regions along the rostro-caudal axis differentially encoded anticipatory threat unpleasantness and its controllability, with FPl serving a uniquely integrative role. First, FIR modeling revealed robust modulation of the amplitude of BOLD responses in LPFC during the anticipation of an unpleasant (*vs.* mild) threat—with FPl coding for threat unpleasantness to a significantly greater degree than mid-LPFC and PMd. This was reflected in both stronger initial responding to unpleasant (*vs.* mild) threat cues, as well as in greater ramping of activation during the anticipation of uncontrollable (*vs.* controllable) unpleasant threats in FPl relative to caudal LPFC regions. Second, representational similarity analyses revealed that threat unpleasantness and controllability information was integrated and maintained in FPl to form putatively high-dimensional conjunctive representations. Notably, the strength of FPl conjunctive representations was associated with better task performance following the anticipation of controllable unpleasant shocks. In contrast, mid-LPFC and PMd held independent representations of threat unpleasantness and controllability that emerged later in the trial. Supporting a potential contribution of amygdala-originated inputs to FPl coding, we found that the magnitude of FPl conjunctive representations was associated with stronger inverse amygdala–FPl functional coupling. Collectively, these results extend prior work on the differential contributions of distinct LPFC regions along the rostro-caudal axis to cognitive control (Badre, 2008; Badre & D’Esposito, 2007; Badre & Nee, 2018; Nee & D’Esposito, 2016, 2017) and align with emerging evidence highlighting the privileged role of FPl in emotion coding (Bramson et al., 2020; Lapate et al., 2022; Roelofs et al., 2023).

A prominent theory of LPFC organization proposes a rostro-caudal hierarchy of cognitive control, in which progressively more rostral regions support increasingly abstract control representations (Badre, 2008; Badre & D’Esposito, 2007, 2009; Koechlin & Jubault, 2006; Koechlin et al., 2003; Koechlin & Summerfield, 2007; Nee & D’Esposito, 2016). These abstract representations are thought to facilitate behavioral control across contexts and timescales (Fuster, 1988; Nee & D’Esposito, 2016). Consistent with this framework, we find evidence for higher-order integration of goal-relevant information in the most rostral region of LPFC—i.e., in FPl. Extending prior work demonstrating mid-LPFC coding of action goals (Lapate et al., 2022), we found that mid-LPFC coded for action controllability independent of unpleasantness. Independent coding of these features was also observed in PMd. While we did not a-priori hypothesize to find evidence of emotion coding in PMd, past work has shown robust recruitment of motor regions when threat stimuli become self-relevant (Portugal et al., 2020).

Conjunctive representations are often hypothesized to be held in a high-dimensional space wherein integrated—and presumably abstract—goal information can be flexibly mapped on to a variety of contexts to promote adaptive behaviors (Badre et al., 2021; Fusi et al., 2016; Kikumoto et al., 2024). Accordingly, we found that the strength of conjunctive threat coding in FPl was associated with greater task accuracy when participants anticipated controllable, unpleasant shocks. Further supporting a high-dimensional, context-specific organizational structure in which FPl representations are embedded, we observed a failure of ‘cross-condition generalization’— whereby an FPl-trained classifier could not generalize task representations (here, threat unpleasantness) across distinct levels of another task factor (i.e., threat controllability) (Bernardi et al., 2020; Bhandari et al., 2024). In contrast, successful cross-condition generalization, together with independent task feature representations (such as held in mid-LPFC and PMd), suggest a lower-dimensional geometry thought to be critical for action selection and execution (Kikumoto et al., 2024; Nee & D’Esposito, 2016). Thus, the present findings extend existing models of LPFC organization and the geometry of cognitive control to naturalistic, emotionally relevant contexts, revealing regionally-specific representational properties along the rostro-caudal axis for the coding of imminent, self-relevant threat and its controllability.

Previous anatomical studies suggest that FPl may have privileged access to amygdala inputs relative to other LPFC regions via the ventral amygdalofugal pathway (Folloni et al., 2019; Ghashghaei et al., 2007; Medalla & Barbas, 2010). Consistently, modulation of BOLD activation by initial threat unpleasantness was strongest in FPl, an effect that progressively attenuated along the rostro-caudal axis. Further evidence for the robust influence of emotional information on FPl function was observed in the progressive ramping of activation immediately prior to the potential delivery of an unpleasant (*vs.* mild) shock, which was greater in FPl than in mid-LPFC and PMd— particularly when the unpleasant threat was uncontrollable. Notably, inverse amygdala–FPl coupling was strongest during the anticipation of uncontrollable unpleasant threats, and stronger inverse amygdala–FPl coupling correlated with stronger conjunctive threat representations in FPl. In contrast, amygdala coupling with mid-LPFC or PMd did not vary as a function of task condition, nor did it correlate with threat-related representations in those caudal LPFC regions.

The robust FPl ramping observed during uncontrollable (*vs*. controllable) unpleasant conditions may be due to inherit differences in uncertainty regarding the final valence and action outcome(s) in each of those trial types, which may in turn modulate amygdala-originated affective signals. In uncontrollable trials, participants are certain that they will receive a shock at the end of each trial, whereas in controllable trials, there is uncertainty regarding the final outcome--i.e., participants do not yet know whether their action will successfully avert the shock, which may result in more ambiguous affective signaling. FPl has been implicated in monitoring and evaluating alternative behavioral strategies and maintaining task-relevant goals (Boorman et al., 2009; Mansouri et al., 2020; Rushworth et al., 2011), and a recent influential theory of emotion control (Roelofs et al., 2023) positions FPl as a key site for dynamically monitoring affective states and guiding flexible responding to emotional challenges (Roelofs et al., 2023). Therefore, FPl may be more engaged in the dynamic exploration of possible courses of action in controllable trials, resulting in more distributed patterns of activity. In contrast, in uncontrollable trials, the unambiguous anticipation of an aversive stimulus may amplify amygdala–FPl threat signaling and contribute to the robust FPl univariate ramping observed here as unavoidable, salient negative outcomes near in time (Bartra et al., 2013; Etkin et al., 2015; Kim & Whalen, 2009; Lapate et al., 2022). Accordingly, as noted above, the uncontrollable unpleasant condition produced the most inverse amygdala-FPl coupling.

Of note, prior research has linked inverse amygdala–PFC coupling to the downregulation of negative emotion (Banks et al., 2007; Herwig et al., 2019; Lee et al., 2012; Morawetz et al., 2017; Salzman & Fusi, 2010; Urry et al., 2006; Wager et al., 2008); thus, it is possible that FPl’s context-sensitive representations may be associated with the down-regulation of amygdala signals that might otherwise drive threat sensitivity and override controllability information. The directionality and functional significance of the inverse amygdala–FPl coupling observed here remain to be fully elucidated. Future work—leveraging combined brain stimulation (e.g., transcranial magnetic stimulation/TMS) and fMRI, for instance—will be required to clarify whether the integrity of FPl function influences the nature of amygdala–FPl coupling during anticipatory threat, and to determine whether bottom up (e.g., amygdala–FPl) versus putatively top-down, FPl-driven amygdala regulatory processes are engaged during processing of threat-relevant information preceding goal-directed action control.

## Limitations

The following limitations of the present work warrant further investigation. First, although the threat-of-shock manipulation we adopted powerfully modulated subjective experience and behavior, it does not permit dissociating whether the coding and integration of emotional information with control signals reported here were driven by negative valence per se, or by a high-arousal state elicited by negatively-valenced stimuli. Future research examining the organization of LPFC in emotional contexts will benefit from including negative, neutral, and positive stimuli to test whether the temporal dynamics and representational properties observed here extend to positively-valenced information, and to disentangle the relative contributions of valence and arousal. In addition, while the fixed countdown to the target-specific action cue and potential shock administration maximized the temporal predictability of threat (Bestmann & Duque, 2016; Cunnington et al., 2003), jittering the interval between the end of the countdown and the motor response would have better enabled disentangling anticipatory emotion and general motor readiness from subsequent action-specific programs, and should be considered in future work. Furthermore, based on participants’ performance, the response window adopted in the current study likely created time pressure and primarily captured neural dynamics related to goal-related urgency. Future work would benefit from varying the motor response window duration to examine how prolonged, strategic control adjustments during response selection and execution influence activation ramping and integrated affect and goal representations in LPFC. Finally, because the associations reported here are correlational, future work employing non-invasive brain stimulation will be essential for assessing whether disrupting FPl signals, including conjunctive affect and control representations at different stages of emotional processing and goal anticipation causally influences behavior and shapes representations in caudal LPFC regions.

## Conclusion

In conclusion, here we demonstrate that FPl—a key node of LPFC circuitry supporting cognitive control—is robustly engaged during the anticipation of emotionally salient experiences and integrates context-sensitive threat and control information. Together, these findings shed light on how emotional inputs shape successful goal pursuit (Shenhav, 2024), echoing Hume’s assertion that “passion” is not an extraneous influence but a central force in guiding action (Hume, 1739).

## Data, Materials, and Code Availability

Analyses were run using custom scripts in FSL, Python and R. Python code used for the Active Escape Task is available on GitHub at https://github.com/LEAPNeuroLab/ASAP/tree/main/Active%20Escape. Data and code used for analysis will be made available on OSF upon publication of the manuscript.

## Acknowledgements

This research was funded by NIH grant MH134000 (R.C.L.) and by Aligning Science Across Parkinson’s ASAP-020-519 through the Michael J. Fox Foundation for Parkinson’s Research (MJFF) (S.T.G. and R.C.L.). The authors thank Taylor Li and Kiana Sabugo for their help with data collection and Mengsi Li for helpful discussions.

## Supplemental Materials

**Figure S1.**
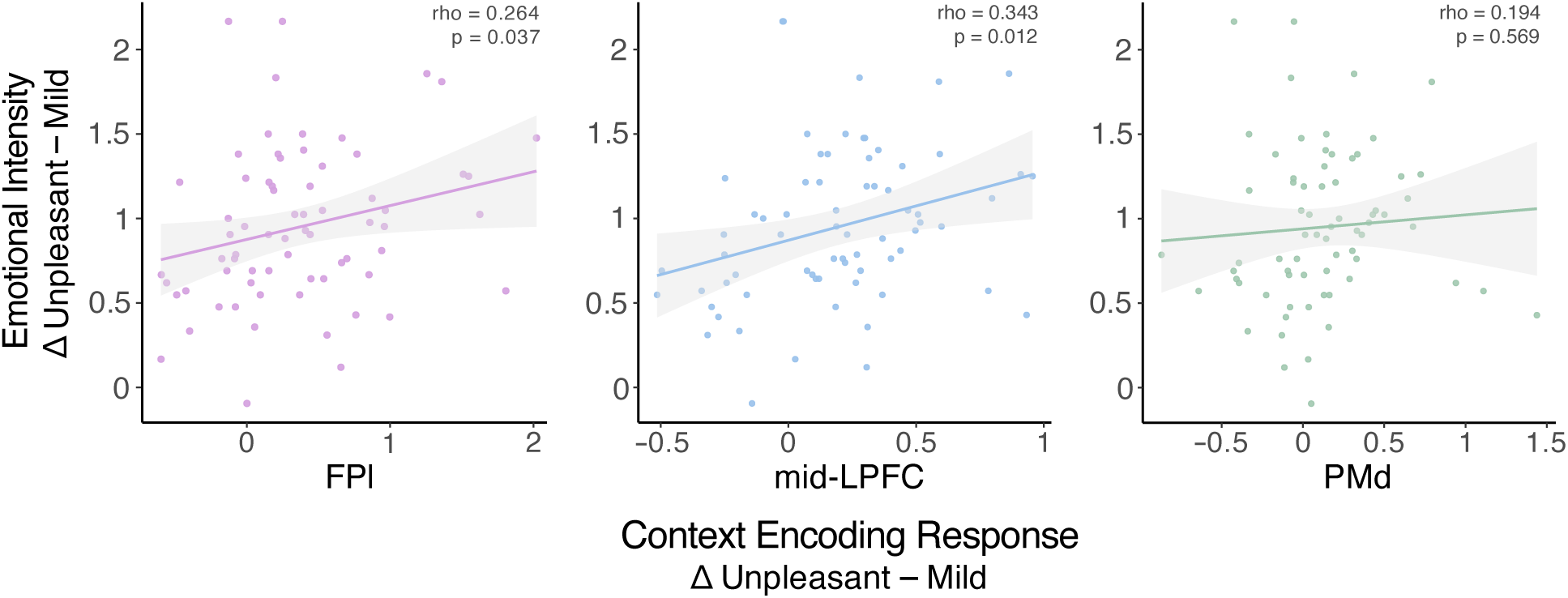
FPl and mid-LPFC initial threat coding correlates with subjective emotional experience. The magnitude of threat unpleasantness-modulated context encoding response (i.e., Unpleasant – Mild) was examined in relation to self-reported emotional intensity to probe the extent to which LPFC coding of threat tracked with subjective experience. A significant positive association between the magnitude of threat coding and self-reported emotional intensity was found in FPl (**left panel**) (*⍴* = 0.25, *p* = 0.04) and mid-LPFC (**center panel)** (*⍴* = 0.30, *p* = 0.01), such that greater initial responding to Unpleasant (*vs.* Mild) shock trials was associated with greater experienced emotional intensity during the countdown period. No association was found in PMd (**right panel**) (*p* = 0.57).

**Figure S2.**
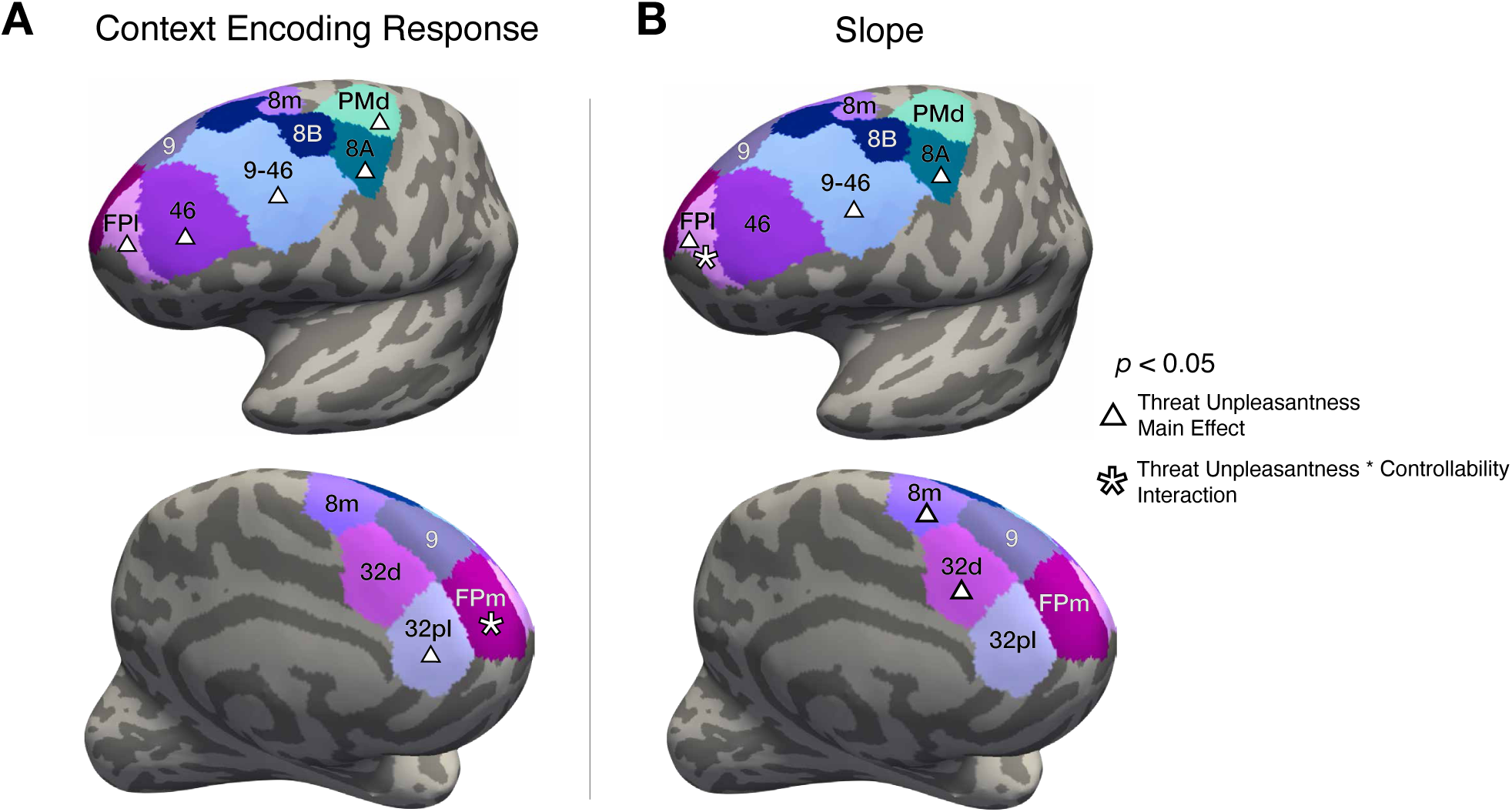
Threat Unpleasantness and Controllability coding in LPFC and mPFC. Significant main effects and interactions of threat unpleasantness and controllability are plotted in regions outside of the a-priori ROIs in LPFC and mPFC separately for the Context Encoding and Shock Anticipation epochs. (**A**) *Context Encoding Response.* Significant main effects of threat unpleasantness and/or significant Threat Unpleasantness * Controllability interactions on the magnitude of the context encoding response are noted (*p* < 0.05). A significant main effect of threat unpleasantness was found in Brodmann’s Area (BA) 8A (*F* = 5.66, *p* = 0.02), BA 32pl (*F* = 4.95, *p* = 0.03), BA 46 (*F* = 35.80, *p* < 0.001), FPl (*F* = 29.04, *p* < 0.001), mid-LPFC [BA 9–46] (*F* = 24.60, *p* < 0.001), and PMd (*F* = 10.33, *p* = 0.001). A significant Threat Unpleasantness * Controllability interaction was found in FPm (*F* = 4.29, *p* = 0.04). (**B**) *Ramping Activation (Slope) during the Shock Anticipation Epoch.* Significant main effects of threat unpleasantness, controllability, and their interaction on the slope of ramping activation during the shock anticipation epoch are noted (*p* < 0.05). A significant main effect of threat unpleasantness was found in BA 8A (*F* = 9.136, *p* = 0.003), BA 8m (*F* = 4.28, *p* = 0.04), BA 32d (*F* = 4.58, *p* = 0.03), mid-LPFC (*F* = 6.59, *p* = 0.01), and FPl (*F* = 4.08, *p* = 0.04). A significant Threat Unpleasantness * Controllability interaction on slope was found in FPl (*F* = 5.19, *p* = 0.02). (Note that results were tested bilaterally but are shown on the left hemisphere for plotting purposes only.)

**Figure S3.**
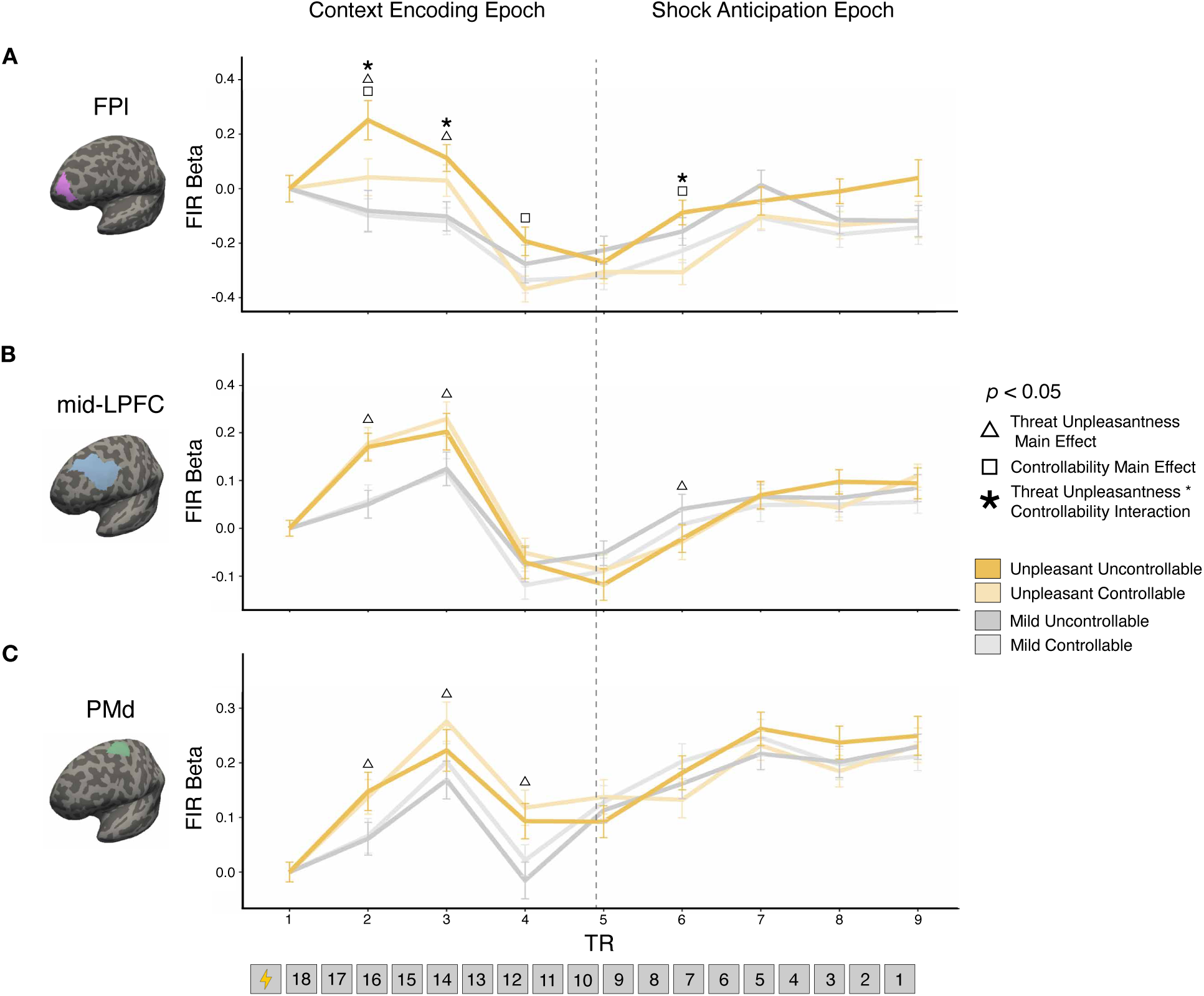
Dynamic changes in information prioritization in LPFC during threat anticipation. Finite impulse response beta coefficients are plotted as a function of threat unpleasantness and controllability in (**A**) FPl, (**B**) mid-LPFC, and (**C**) PMd. Anticipatory threat unpleasantness and controllability significantly modulated (**A**) FPl function over time [controllability main effect: F = 23.36, *p* < 0.001; threat unpleasantness main effect: *F* = 8.48, *p* = 0.004; Threat Unpleasantness * TR interaction: *F* = 3.27, *p* < 0.001]. A two-way Threat Unpleasantness * Controllability interaction in FPl (*F* = 5.19, *p* = 0.02) was driven by significantly stronger FPl responding to Unpleasant *vs.* Mild trials in the Uncontrollable condition [Uncontrollable Unpleasant – Mild: *B* = 0.09 (SE = 0.02), *t* = 3.67, *p* = 0.001, *d_z_* = 0.21, 95% CI [0.03, 0.15]] relative to the Controllable condition [Controllable Unpleasant – Mild: *p* = 0.97]. Threat unpleasantness significantly modulated activation in (**B**) mid-LPFC [threat unpleasantness main effect: *F* = 4.01, *p* < 0.05; Threat Unpleasantness * TR interaction: *F* = 3.68, *p* < 0.001] and (**C**) PMd [threat unpleasantness main effect: *F* = 7.95, *p* = 0.005; Threat Unpleasantness * TR interaction: *F* = 2.26, *p* = 0.02]. No main effects of controllability (all *p*s > 0.26), nor Threat Unpleasantness * Controllability interactions were found in mid-LPFC or PMd (all *p*s > 0.43). Consistent with the finding that only FPl showed a significant Threat Unpleasantness * Controllability interaction, the overall Region * Threat Unpleasantness * Controllability interaction was also significant (*F* = 3.13, *p* = 0.04). Error bars denote the within-subjects 95% confidence interval for each condition, computed using Morey’s (2008) method (Morey, 2008).

**Figure S4.**
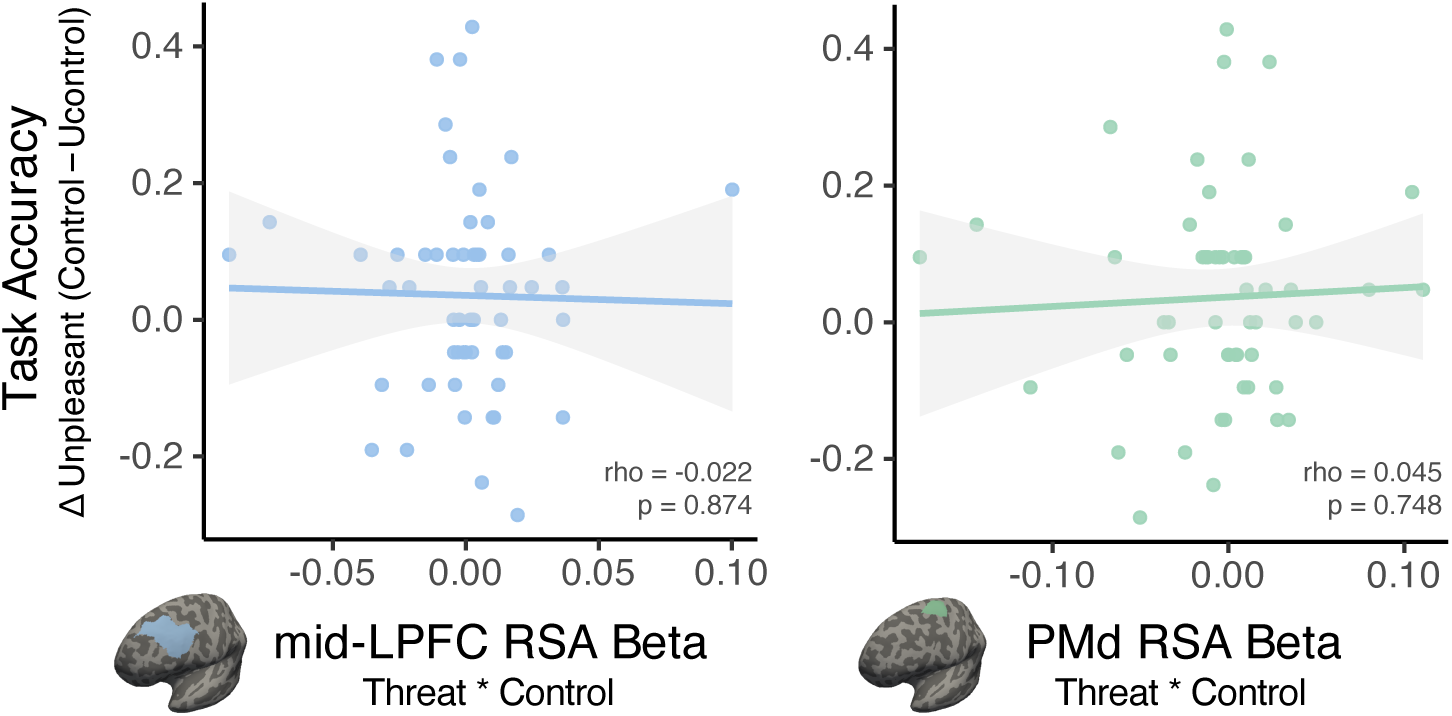
Conjunctive coding of threat and controllability is not associated with task accuracy in mid-LPFC or PMd. Conjunctive Threat Unpleasantness and Controllability betas did not correlate with task performance in Controllable Unpleasant (*vs.* Unpleasant) in mid-LPFC (*⍴* = −0.02, *p* = 0.87; **left panel**) or PMd (*⍴* = 0.05, *p* = 0.75; **right panel**).

